# Position-theta-phase model of hippocampal place cell activity applied to quantification of running speed modulation of firing rate

**DOI:** 10.1101/714105

**Authors:** Kathryn McClain, David Tingley, David Heeger, György Buzsáki

## Abstract

Spiking activity of place cells in the hippocampus encodes the animal’s position as it moves through an environment. Within a cell’s place field, both the firing rate and the phase of spiking in the local theta oscillation contain spatial information. We propose a position-theta-phase (PTP) model that captures the simultaneous expression of the firing-rate code and theta-phase code in place cell spiking. This model parametrically characterizes place fields to compare across cells, time and condition, generates realistic place cell simulation data, and conceptualizes a framework for principled hypothesis testing to identify additional features of place cell activity. We use the PTP model to assess the effect of running speed in place cell data recorded from rats running on linear tracks. For the majority of place fields we do not find evidence for speed modulation of the firing rate. For a small subset of place fields, we find firing rates significantly increase or decrease with speed. We use the PTP model to compare candidate mechanisms of speed modulation in significantly modulated fields, and determine that speed acts as a gain control on the magnitude of firing rate. Our model provides a tool that connects rigorous analysis with a computational framework for understanding place cell activity.

**Significance:** The hippocampus is heavily studied in the context of spatial navigation, and the format of spatial information in hippocampus is multifaceted and complex. Furthermore, the hippocampus is also thought to contain information about other important aspects of behavior such as running speed, though there is not agreement on the nature and magnitude of their effect. To understand how all of these variables are simultaneously represented and used to guide behavior, a theoretical framework is needed that can be directly applied to the data we record. We present a model that captures well-established spatial-encoding features of hippocampal activity and provides the opportunity to identify and incorporate novel features for our collective understanding.

## Introduction

Place cells in the rodent hippocampus encode spatial information through their spiking activity. As the rodent moves through an environment, the firing rate of place cells increases at particular locations, termed “place fields”, suggesting a firing rate code for position (1). Meanwhile the local field potential in the hippocampus is dominated by a 7-9 Hz ‘theta’ oscillation and place cell spiking is modulated according to the phase of this oscillation (2). The phase at which spiking occurs precesses as the animal moves through the place field, a phenomena known as “phase precession” (2). Two overlapping codes for position emerge: a rate code and a phase code.

It has been suggested that the phase code is used to identify the animal’s location, while the firing rate can be used to encode other variables, such as the speed of the animal’s movement (3). Indeed, firing rate variability from trial to trial has been a long noted and unexplained feature of place cell activity (4-6). In support of this hypothesis, several papers have reported a positive correlation between running speed and firing rate of pyramidal neurons in the hippocampus (7-9), entorhinal cortex (10-13), and neocortical neurons (14, 15). Replicability of this effect in hippocampus has only recently been questioned (16).

Analyzing the influence of additional variables (such as running speed) on place cell firing is difficult for several practical reasons. First, the experimenter has only limited control over the relevant variables (position, running speed, theta phase), making the behavioral paradigm nearly impossible to design without introducing additional interfering elements. Second, rodents will only run a small number of trials on a given day, typically dozens, giving us limited statistical power for analyzing effects across multiple dimensions.

Another challenge in understanding this system is the dynamic interaction between the rate code and phase code. As these codes combine, different formats of information are conveyed simultaneously in place cell spiking. The interaction can produce unintuitive, though entirely predicable results in traditional analyses of place cell activity. These practical challenges have hindered the effort to explain variability in hippocampal firing rates. A computational tool is needed that accounts for the well-established features of this system and provides a path forward in asking further questions.

We have drawn from classic GLM (13, 17), gain control models (18-20) and phase-precession models (21, 22) to develop a position-theta-phase (PTP) model of place cell activity. In this model, spatial input is scaled by theta phase modulation to determine the firing rate of a place cell. This model has three primary utilities:

1. It provides a quantitative description of the relevant features of place cell activity. The model can be reliably fit with fewer than 100 spikes. These descriptive statistics can be compared across time, conditions and cells.
2. The model can easily generate simulated place cell data that mimics real place cell activity. The simulated data can be used for analysis on its own, or as statistical grounding in analyses of real data.
3. It introduces a framework for principled hypothesis testing. By iteratively adjusting features of the model and using a model fit comparison with the proposed baseline model, we can assess the influence of additional modulating variables.

We demonstrate these utilities and use the model to assess speed modulation in place cells. We find no evidence for speed modulation in the majority of place fields. For a minority of place fields, spiking is either positively or negatively modulated by speed. The modulation appears to act as a gain control on the overall magnitude of firing, as opposed to other candidate computations. Our PTP model offers a disciplined tool to separate physiological mechanisms from spurious statistical artifacts that may result from non-intuitive interactions of observed variables.

## Results

### Parametric model of place cell activity

We model place cell activity as a function of two independent inputs: position and theta phase (Figure 1A). Place cell activity is canonically described as a Gaussian function of position over select portions of the environment. These portions (“place fields”) are analogous to sensory receptive fields for place cells. Place cell spiking outside of the place field is typically sparse and has not been characterized, therefore our model focuses on within-place-field spiking. The spatially-activated responses are modulated according to the phase of the theta oscillation, and the preferred phase of the modulation changes with position, precessing to earlier phases of the theta cycle as the animal moves through the field (2, 22). This endows the phase of each spike with spatial information, along with the magnitude of the firing rate. We formalized these concepts by modeling firing rate as a Gaussian spatial response function, scaled by a Von Mises theta modulation function (Figure 1B). The preferred phase of the theta modulation function shifts with position according to the linear precession function. We modeled spiking as an inhomogeneous Poisson process of the firing rate.

**Figure 1:**
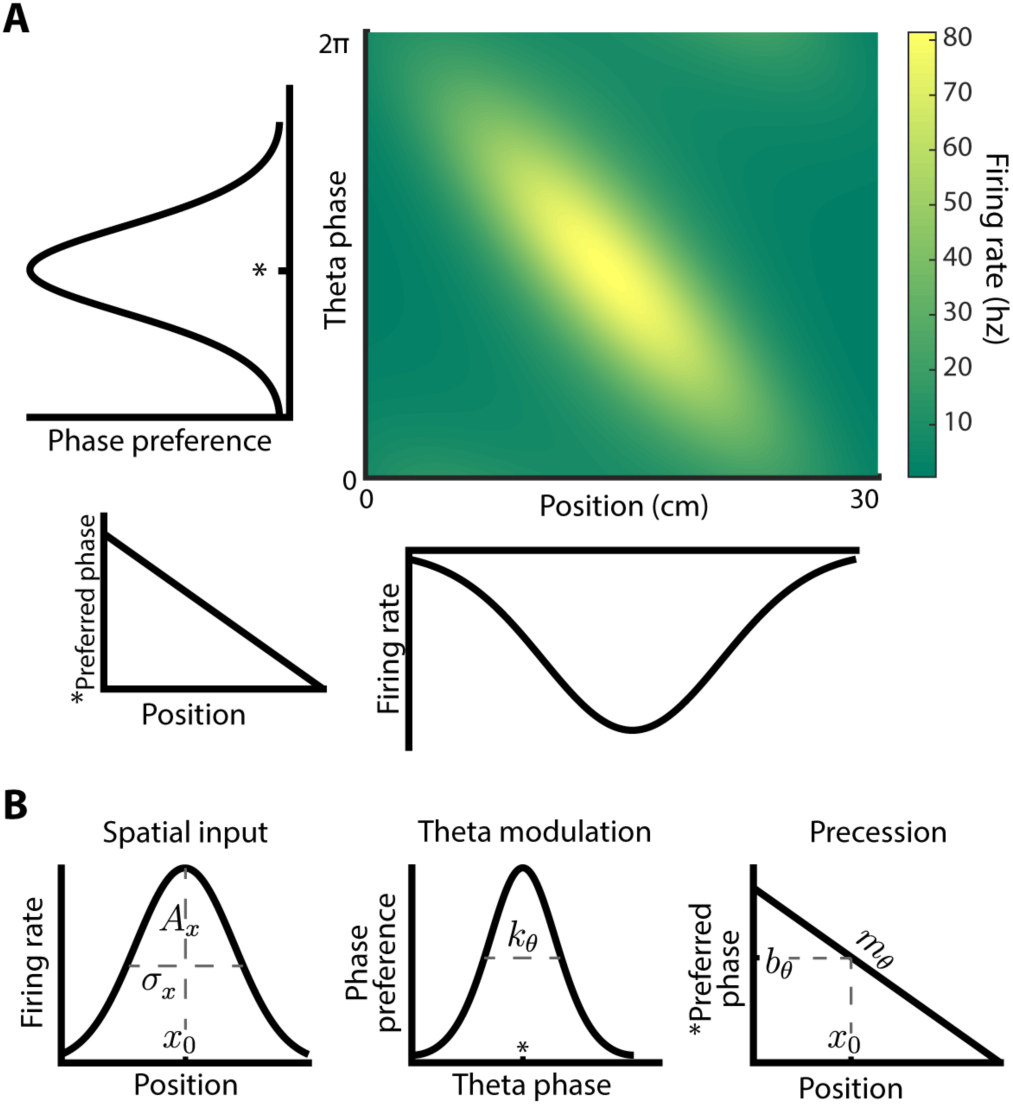
Parametric model of place cell activity. A) Schematic of model: firing rate modeled as a multiplication of tuning curves with respect to space and theta phase. Preferred theta phase changes with position to incorporate phase precession. Parameters for this schematic derived from fitting model to real place field data. B) Model equations: Spatial input function is a Gaussian function of position with 3 parameters: amplitude *A*_*x*_, width *σ*_*x*_ and center *x*_0_. Theta modulation function is a Von Mises function (approximately circular Gaussian) of theta phase normalized to height 1 with one parameter: *k*_*θ*_. Theta modulation is centered on a preferred phase in the precession function which changes linearly with position according to slope *m*_*θ*_ and intercept *b*_*θ*_.

The PTP model functions are determined by parameters that correspond to relevant functional features of place cell activity (Figure 1B). The amplitude of the firing rate is parameterized by *A*_*x*_. The width and position of the place field are captured by *σ*_*x*_ and *x*_0_, respectively. The theta-phase selectivity is determined by *k*_*θ*_. Cells that spike within a narrow range of phases have high selectivity while cells that spike across the whole cycle have low selectivity. *m*_*θ*_ determines the rate of phase precession and the preferred phase at the center of the place field is *b*_*θ*_.

### Fitting and simulating place cell activity

We fit the model to spiking data recorded from place cells in rats. We examined data recorded from dorsal CA1 region of the hippocampus in rats as they ran along linear tracks (23). We used the rat’s position, theta phase and spike timing to estimate parameter values for each place field (see methods; Figure S1). Parameter values were stable across random subsets of data (Figure S2), indicating that our fitting procedure is reliable for this quantity of experimental data. With the parameter estimates for individual place fields as well as the distribution of parameters across the whole population, we were able to generate empirically grounded simulation data.

Using parameters estimated from an example place field, we generated place cell spiking in a simulated experiment (Figure 2). For each simulated trial a virtual rat ran through the place field at a constant speed while theta oscillated at a constant frequency, resulting in a straight trajectory through phase-position space (Figure 2Ai). The PTP model predicted the firing rate at each point along this trajectory and simulated spiking as a stochastic Poisson process of the instantaneous firing rate (Figure 2Aii). From one trial to the next, the initial theta phase at the beginning of the field randomly shifted, as it does in real experiments, which resulted in spatially shifted firing rate patterns.

**Figure 2:**
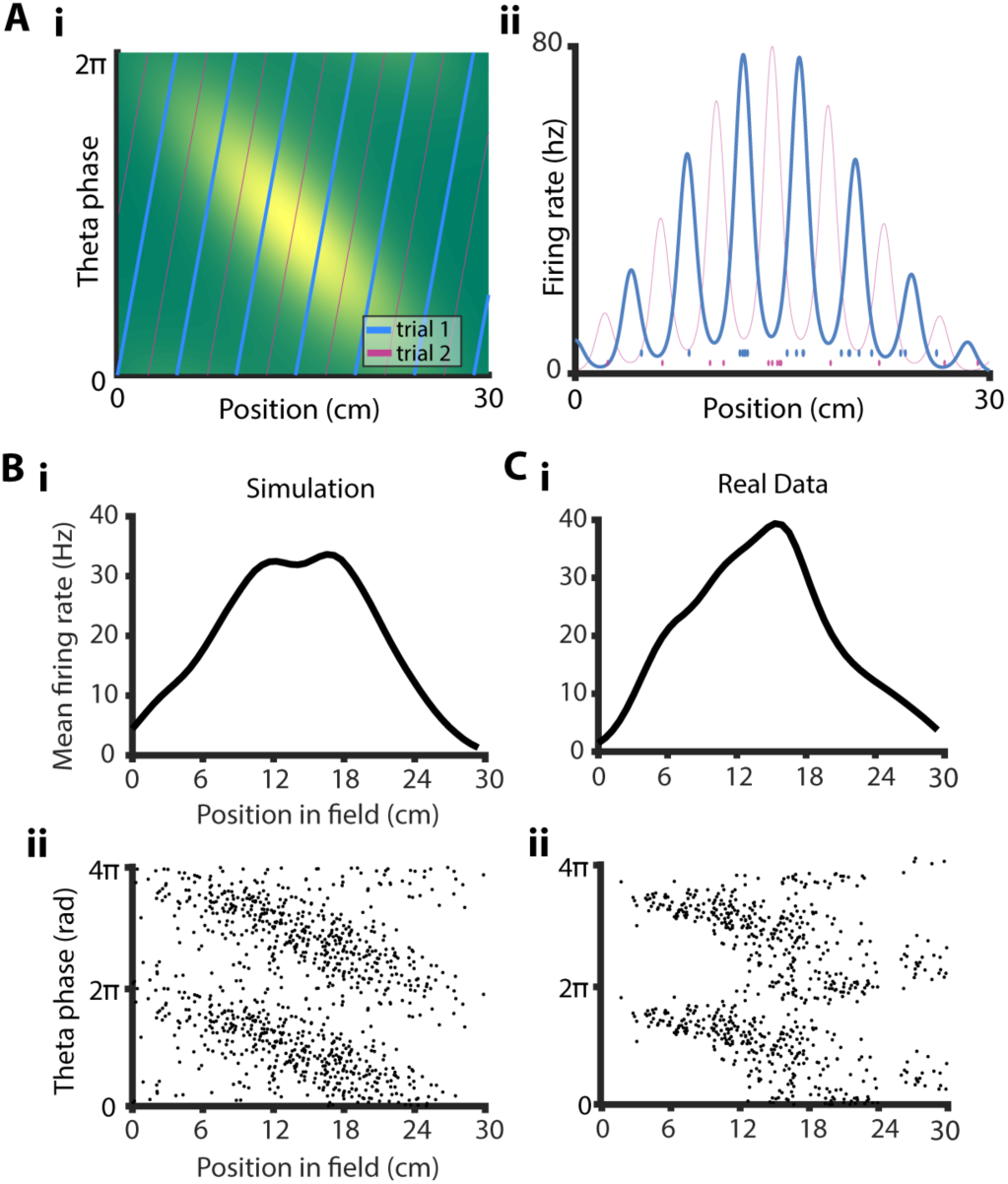
Simulating place cell data. A) Single trial simulation

i. Simulated theta-phase-position trajectory imposed on place field schematic (as in Figure 1A) for trial 1 (blue), and trial 2 (purple) at the same speed with a different initial theta phase at the start of the place field.
ii. Simulated firing rates for model fit to place field in C) computed for trajectories in trial 1 and 2 in i. Spiking for each trial (below) simulated via Poisson process. B) Summary visualization of simulated data: Spiking was simulated for 42 trials with varying speeds and initial theta phases, using the PTP model fit got real place cell data in C).

i. Mean firing rate vs. position, averaged over trials.
ii. Theta phase vs. position for each spike. C) Summary visualization of real place cell data in example place field. i. and ii. same as in B).

The magnitude of firing rates predicted by the PTP model is larger than typically reported for place cells. The range of firing rates for place cells within their place field has been reported as 1-40 Hz based on the trial-averaged firing rate within the place field (3, 24). However, on individual trials, our model predicts 80-100 Hz peak firing rate for many cells (Figure 2Aii, Figure S4). The discrepancy arises from trial averaging, which averages out the effect of the theta modulation, resulting in an averaged peak firing rate roughly half the true peak firing rate (Figure 2Bi)

Inter-trial randomness in the theta phase at the beginning of the place field could also explain part of the variability in place cell firing rates. We simulated two trials with the same parameters and running speed, varying only the initial theta phase at the entry to the place field (Figure 3A). The theta modulation was shifted with respect to the spatial input, which changed the predicted firing rate across the field by a factor of two. Including the Poisson variability of spiking, the firing rate within the field could reasonably be 4 Hz on trial one and 16 Hz on trial two. This difference is based solely on the initial theta phase and Poisson variability.

**Figure 3:**
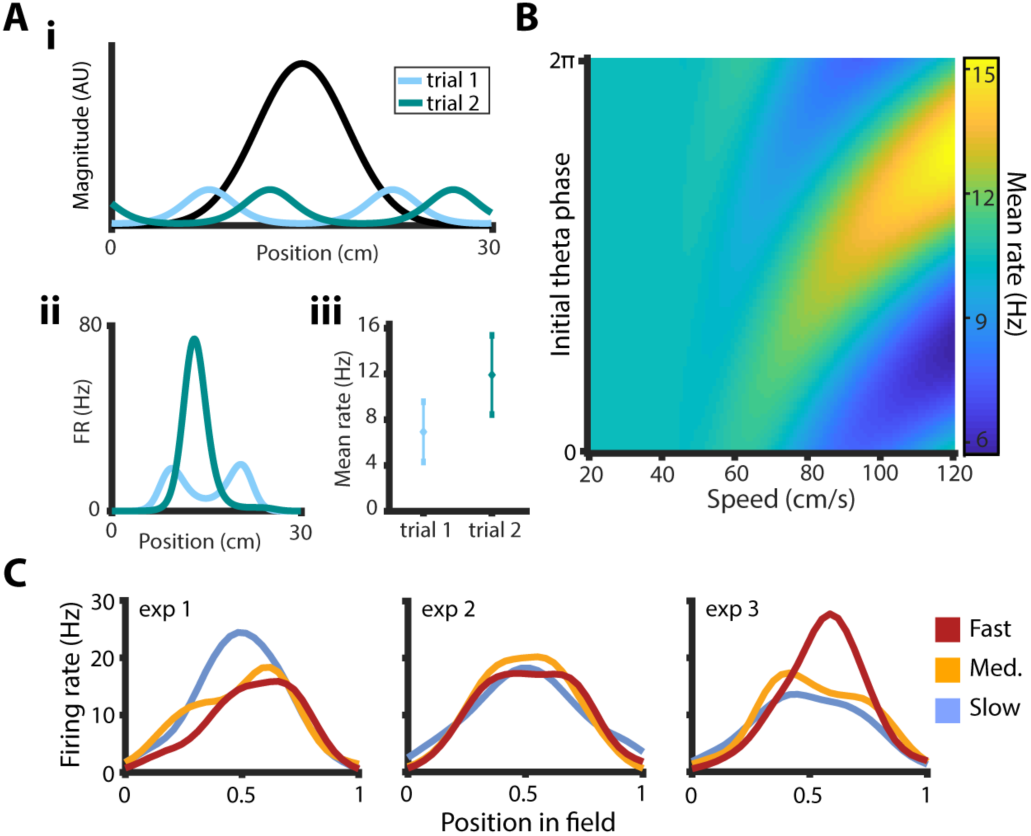
Speed-dependent variability can cause spurious correlations with running speed. A) Two simulated trials with identical model parameters and running speed, differing only in initial theta phase.

i. Alignment between spatial input function (black) and phase modulation function for trials 1 and 2 (blue and green).
ii. Predicted firing rate vs. position for trials 1 and 2 computed by multiplying the spatial input and phase modulation functions in i.
iii. Mean firing rates for trials 1 and 2, averaged over position. Error bars correspond to standard error (SEM) predicted from Poisson variance of spiking. B) Mean firing rate across place field simulated as a function of initial theta phase and running speed. Variability caused by initial theta phase increases at higher speeds. C) Firing rate vs. position in the fast (red), medium (yellow), and slow (blue) sets of trials in three simulated experiments. In each experiment 30 trials were simulated with randomized speeds and initial theta phases. Mean firing rate was computed as a function of position for each set of trials. Conditions were identical for each simulated experiment with no explicit speed dependence in the model, however apparent speed modulation appeared by chance, both negatively (experiment 1) and positively (experiment 3).

The effect of the initial theta phase on firing rate is amplified at faster running speeds. We used the PTP model to predict the firing rate as a function of both running speed and initial theta phase (Figure 3B). At slow speeds the initial theta phase is not very influential in the overall rate; however, at fast speeds the expected rate can vary dramatically. The intuition behind this observation is when the animal runs slowly, many theta cycles occur within the place field, making the alignment of any particular cycle less important for the overall predicted firing rate. At faster speeds, there are fewer cycles, making the coincidence of the theta modulation and spatial input much more important.

The speed-dependent variability could produce spurious correlations between speed and firing rate. We simulated a place cell experiment three times with identical conditions (Figure 3C). In each experiment we randomly varied running speed and initial theta phase, drawing from a uniform distribution of each, and used the model to generate spikes. We computed the average firing rate at each position for the fast, medium and slow trials. Across the three simulations, an apparently negative relationship between speed and firing rate arose in one, no relationship was evident in another, and a positive relationship appeared in the last. Recall the model used for simulation has no explicit speed dependence, so each of the apparent relationships is artifactual. The confound between running speed and firing-rate variability makes the analysis of speed tuning in place cells difficult, because standard statistics are not sufficient to assess the significance of these relationships.

### Quantifying the effect of running speed on place cell activity

In real place field data, we found a heterogeneous distribution of speed dependence using standard correlations. We computed the average speed and firing rate within the place field for each trial and calculated the correlation across trials. Contrary to previous findings that have reported mostly positive correlations between running speed and firing rate (3, 7), and in line with a more recent report (16) we found place fields with ostensible negative speed relationships (Figure 4A) as well as those with apparently positive relationships (Figure 4C). We also found a majority of place fields that did not appear to be speed modulated (Figure 4B). Across all place fields in the dataset, the distribution of correlations appeared to be heterogeneous (Figure 4D). However, because speed related variability can produce spurious correlations (Figure 3C) these apparent effects must be scrutinized.

**Figure 4:**
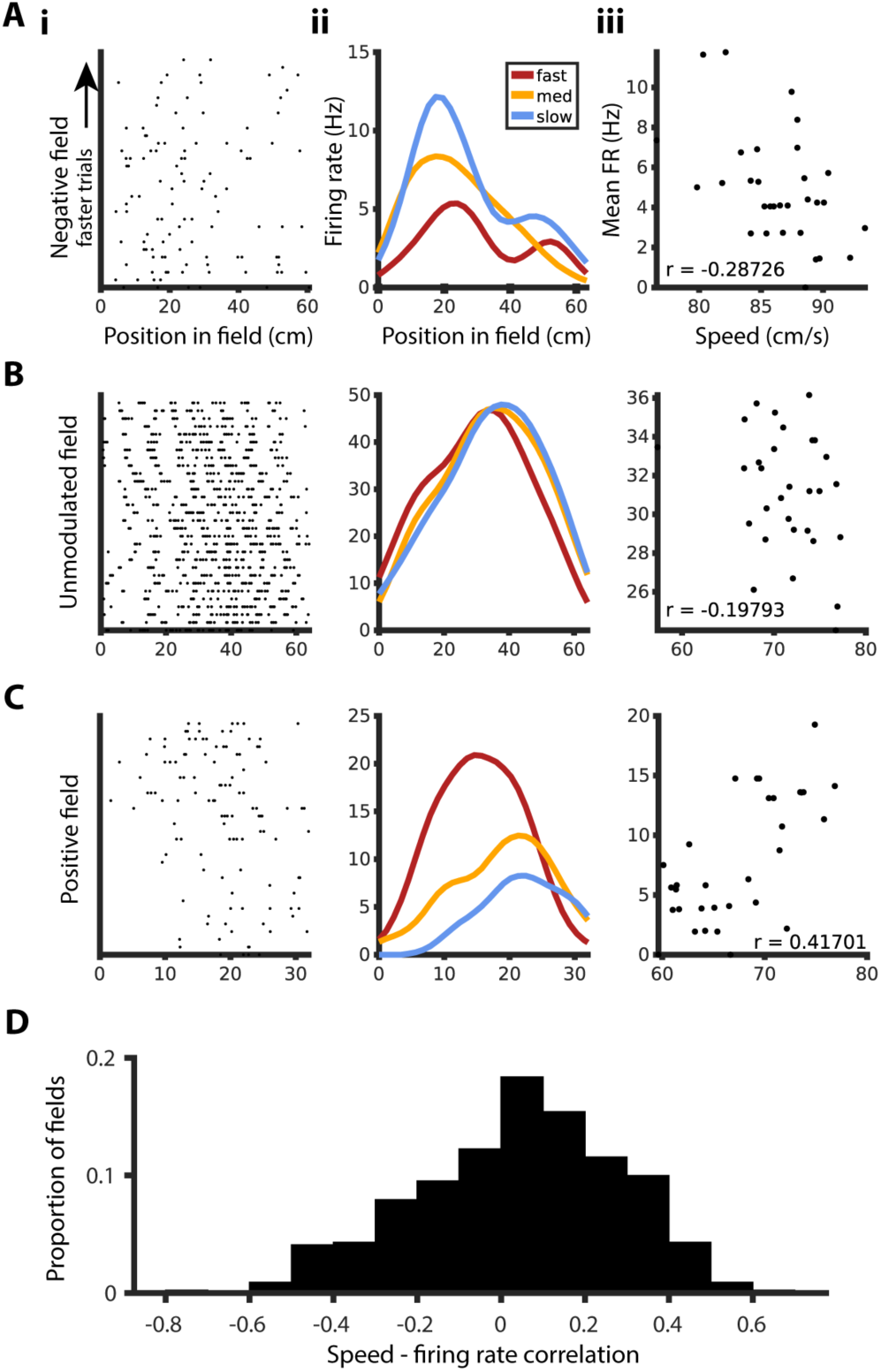
Real pyramidal cells show heterogeneous distribution of speed correlations. A) Speed dependence of example place field with negative speed modulation:

i. Trial vs. position of each spike, trials ordered by mean running speed in the place field.
ii. Firing rate vs. position for fastest (red), middle (yellow) and slowest (blue) thirds of trials.
iii. Mean firing rate across place field vs. speed, each point representing one trial. r values throughout indicate Kendall rank correlation coefficient. B) Same as A) for unmodulated example place field. C) Same as A) for positively speed-modulated place field. D) Distribution of speed-firing rate correlations across all place fields in dataset.

To more rigorously assess speed modulation, we used the PTP model to generate an ensemble of simulated experiments, from which we computed a null distribution of speed correlations. We use “null distribution” because the PTP model has no speed dependence, so the resulting correlations arise solely from the sources of variability accounted for in this model. For each place field in our dataset, we used the estimated model parameters to virtually recreate the experiment (Figure 5A). We computed the correlation between speed and simulated firing rate, then repeated the simulated experiments 20,000 times to compute a null distribution of correlation values (Figure 5B). The null distribution varies across place fields depending on the best-fit model parameters for each individual field, and so must be computed independently.

**Figure 5:**
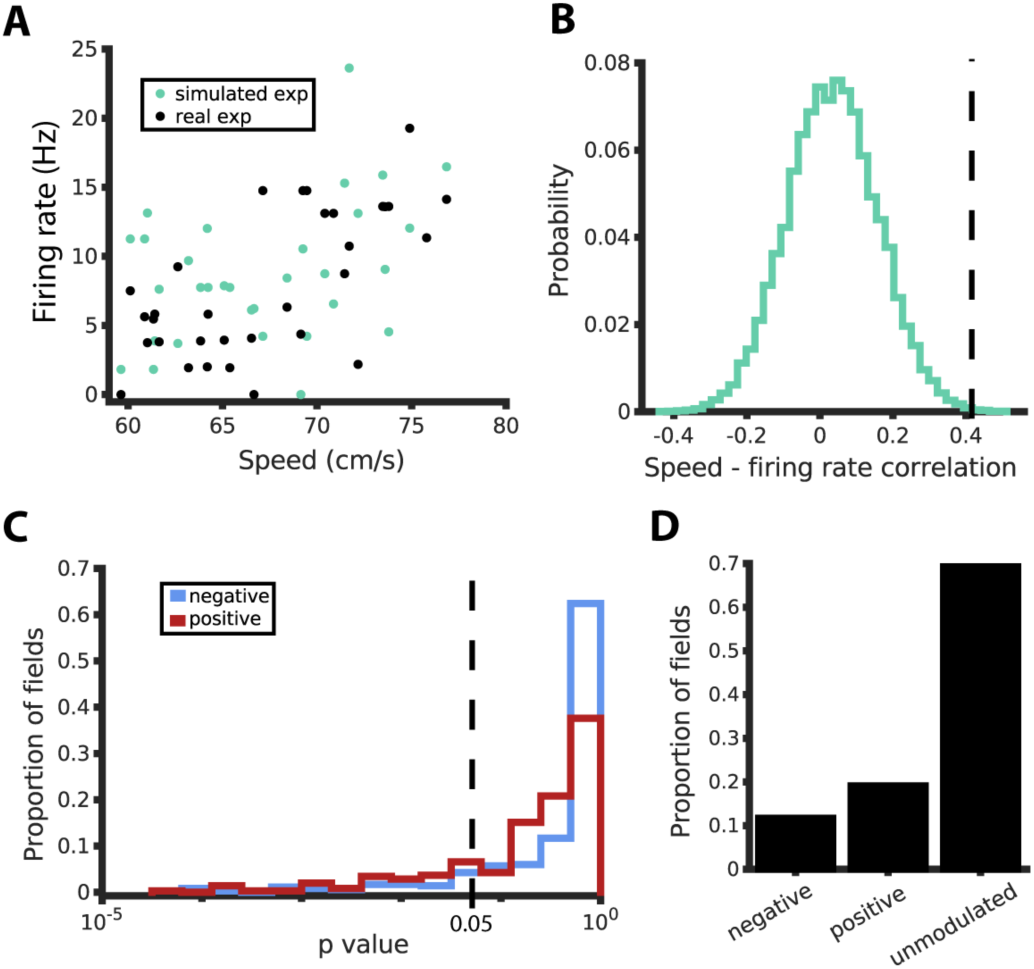
Speed modulation is statistically significant in a small number of place fields. A) Mean firing rate vs. speed for real example place field (same as Figure 4C), for real experiment (black) and simulated experiment (green). Simulation performed using model fit to this place field and conditions identical to real experiment. B) Null distribution of speed-firing rate correlations (green) computed from 20k simulations of the experiment in A). Empirical correlation in black. For this example field, we find evidence for speed modulation beyond what can be explained by the PTP model. C) One-tailed p-values computed from null distribution for all place fields in the dataset. Significant speed modulation is defined as p<0.05. Red, distribution for positive speed-firing rate relationships. Blue, distribution for negative speed-firing rate relationships. D) Proportion of place fields for which there was statistical significance for negative or positive speed modulation, and the proportion for which there was no evidence for speed modulation.

Speed-firing rate correlations for most place fields did not lie significantly outside the respective null distributions. The distance between the true correlation value and the null distribution was measured as a p-value in the positive and negative directions (Figure 5C). A criterion of p<.05 was used and fields with significant positive and negative modulation were identified. A subset of place fields with positive modulation (19%) and a subset with negative modulation (12%) were identified above chance levels. Nonetheless, the majority of place fields (69%) did not meet our criterion for significant speed modulation (Figure 5D). In summary, we found that the degree of speed modulation in the majority of place fields lies within the predictions of the PTP model, which does not include any speed dependence. Yet, we found a minority of place fields with speed modulation beyond the model predictions. Next, we sought the potential mechanisms of such effects.

We extended the PTP model to explore the computational effect of speed on firing rates of significantly modulated place fields. The model delineates the independent features of place fields that could be affected by running speed. Two candidates that could directly impact the average firing rate in the place field are the amplitude *A*_*x*_ (Figure 6A), and the phase selectivity *k*_*θ*_ (Figure 6B). Modeling amplitude as a function of speed corresponds to a gain control model, where running speed multiplicatively scales the magnitude of activity. Modeling phase selectivity as a function of speed corresponds to a changing window of spiking within the theta cycle. These two mechanisms could also work in consort in a dual speed model (Figure 6C). In each of these speed-dependent variants of our original model, we model the speed-dependent parameter as a linear function of speed. The slope of this function determines the direction and degree of speed dependence.

**Figure 6:**
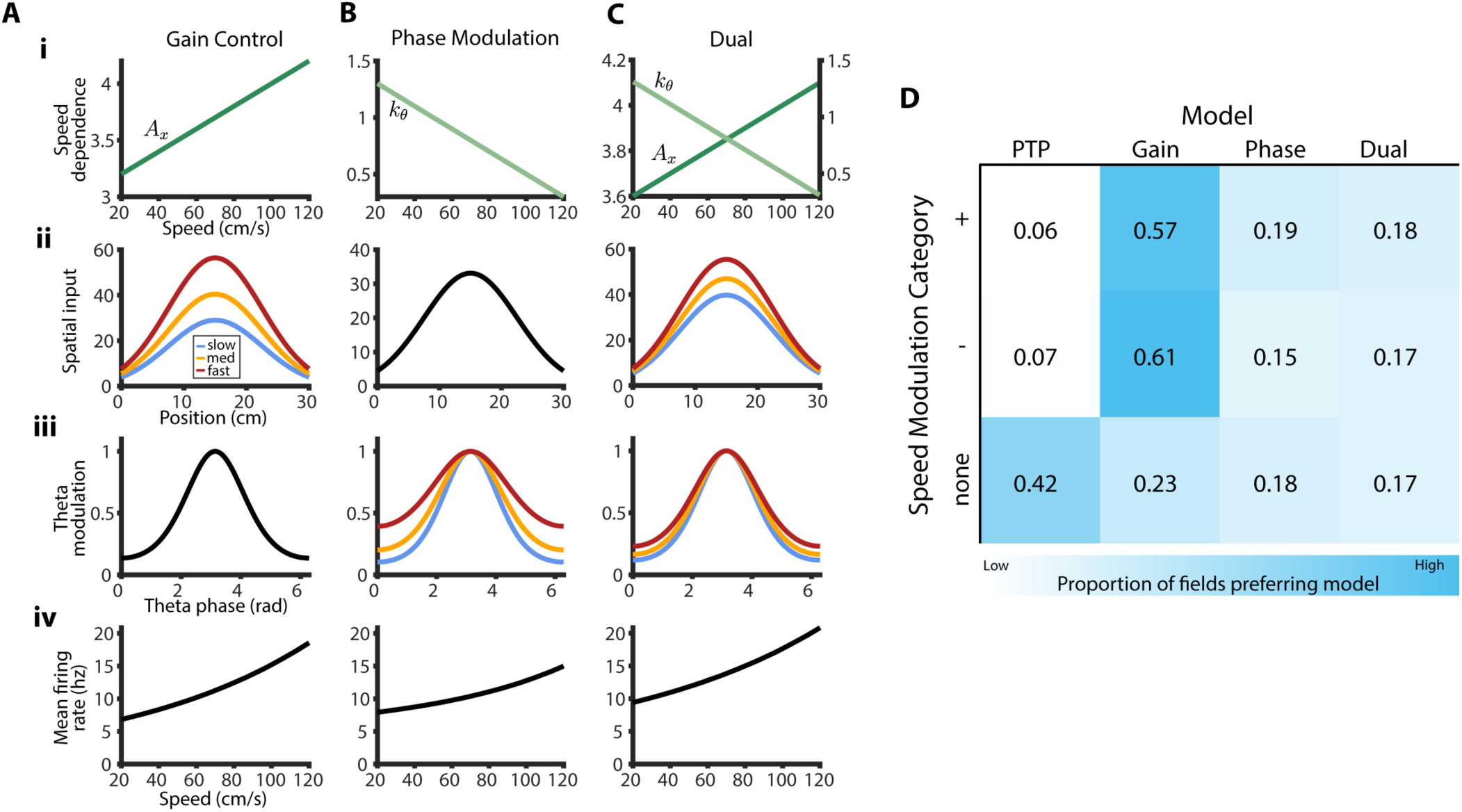
Speed model comparison: variants of basic model that include speed dependence. A) Gain control model:

i. Hypothetical relationship between amplitude parameter and speed.
ii. Spatial input function as it varies with speed.
iii. Phase modulation function (stationary with respect to speed).
iv. Hypothetical relationship between firing rate and speed for gain control model. B) Phase modulation model: same as A) except phase selectivity varies with speed instead of amplitude. C) Dual modulation model: same as A) except both amplitude and phase selectivity vary with speed. D) Proportion of place fields in each speed modulation category best fit by each model variant. Positively and negatively modulated fields are by majority best fit with a gain control model, while unmodulated fields are mostly best fit with the original PTP model.

Significantly speed-modulated place fields are best explained by a gain control model of speed. We performed a model fit comparison for each place field, comparing the speed-dependent model variants and the original place field model. For each place field we fit each model and measured the performance of the model in predicting a held-out subset of the data. We selected the model with the highest average log-likelihood as the preferred model for that place field (see Methods). Expectedly, the majority of unmodulated place fields preferred the original PTP model (Figure 6D). Among the subsets of positively and negatively modulated place fields, the majority preferred a gain control model of speed modulation. These results suggest the computational effect of speed operates primarily on the magnitude of place cell activity, leaving the theta-phase modulation of place cells unaffected.

## Discussion

### Refining previous notions of speed modulation of firing rates

Our analysis of speed modulation in place cell activity provides some amendments to previous notions in the spatial navigation field. We did not find evidence for speed modulation in the majority of place cells and suggest increased firing rate variability at high speeds as a potential source of spurious correlations. Of the minority of fields that did show modulation, some were positively modulated and others were negatively modulated. For each of these subsets, speed appears to affect the activity primarily as a gain control, scaling the overall magnitude while theta modulation remains mostly unaffected.

The lack of robust speed dependence of place cell firing rates may convey an important robustness of the system. If place cells are used to for navigation purposes, altering the ‘code’ for behavioral parameters such as running speed may not be advantageous. Interleaving codes through gain control computations can allow one population to simultaneously represent multiple variables (3, 19, 25-27), and our results do suggest that speed dependence in a minority of place fields is best characterized as gain control. However, the sparsity of place cells in our data showing any speed dependence makes this interpretation tenuous.

An additional consideration in studying “speed modulation” is the relationship between speed and trial number that exists in almost all experiments. As the animal’s motivation decreases throughout the course of an experiment, running speed also decreases, causing an inseparable correlation between speed and trial number (Figure S5). This may be a confounding variable as neural signals associated with velocity are reciprocally woven into neural circuits that control motivated behavior (28-30). What has been identified as “speed modulation” in this report, and likely in others, could also be considered a motivation signal modulating activity, or simply “drift,” *i.e.* slow changes in activity patterns over time. In terms of functionally characterizing sources of variability in the system, such a distinction may not be important, because speed, time and motivation are correlated. However, if the goal is to identify underlying physiological mechanisms of the effect, it should become an important consideration.

A quantitative characterization of drift over time in place field activity is a much-needed analysis for hippocampal research that our PTP model would be suited to address. As experimenters probe physiological circuits by performing manipulations and recording multiple changes, a baseline characterization of the volatility is needed to specify the effects caused by the manipulations and separate them from appealing, though ultimately spurious effects.

Our findings do not contradict suggestions that speed is a fundamental parameter of hippocampal activity. We found that the firing rates of the majority of putative fast-spiking interneurons, but not those of slow-spiking interneurons, were positively modulated by running speed (Figure S6). Fast-spiking interneurons, rather than pyramidal cells or slow-spiking interneurons, may be responsible for speed control of frequency of theta oscillation of hippocampal place cells (2, 21, 31, 32) and entorhinal grid cells (33).

### The PTP model: uses and findings

The PTP model we describe here provides a functional description of the well-established factors that influence place cell activity: position and theta phase. Position is an external variable that exists in space, while the theta oscillation is entirely internally generated and propagates in time. These variables interact dynamically through running speed, which may exert its own place-field-specific influence on activity. The results of this interaction are not always obvious or intuitive. Our model can be used in lieu of intuition to inform baseline controls. Appropriate controls are necessary to ward against interpreting inherent implications of the position-phase interaction as novel features of place cell activity. Our model also provides a framework for identifying and incorporating truly novel features into our collective understanding of hippocampal operations.

The PTP model has allowed us to uncover a few surprising features of place cell activity. First, the dynamic range of place cell firing rate is roughly double what is typically measured from trial-averaged firing rates. Second, running speed affects firing rate variability due to Poisson randomness and alignment between theta phase and position, which can produce spurious correlations between speed and firing rate. Finally, despite the potential for spurious correlations, there appear to be small subsets of place fields that show genuine speed modulation.

In general, the PTP model can be used in several ways. First, key features of place fields can be described quantitatively by fitting the model with relatively small amounts of place cell data. Second, realistic place cell data can be generated in simulated experiments, with conditions and parameters fully controlled by the experimenter. Simulated place cell data can help explore theoretical aspects of the hippocampal spatial navigation system, inform the design of future experiments and serve as a control in analyzing real place cell data. Lastly, hypothetical variations of the model can be systematically tested to uncover additional features of the data as demonstrated in Figure 6.

### Model limitations and extensions

In formulating our model, choices were made for the sake of simplicity that may have neglected some particular specifics of hippocampal physiology. For example, formulating phase precession as a linear function of position (2, 3) ignores previous work that has characterized a curved “banana” shaped phase precession (24, 34). However, the PTP model can be amended to accommodate details of such properties, and be a useful tool for further probing their significance.

Physical stationarity of the place field is a more fundamental assumption of the PTP model. We define the spatial input function as an environmental input drive at a particular location (1). An alternate interpretation is that a place field begins at the occurrence of the first spike, and the place field peak varies from trial to trial (5, 6), tying the place field more strongly to theta phase than to position. This interpretation may be useful in some contexts, but ours reflects common assumptions in the field that are arguably more relevant in the context of spatial navigation.

One-dimensional space is another assumption. In its current instantiation, the PTP model is not directly applicable for two-dimensional navigation. However, with additional assumptions it could be expanded to two dimensions.

We also only model single place fields, while real place cells can have multiple fields within an environment (35, 36). As is, PTP models for multiple fields could easily be combined along an expanded position axis. A potentially interesting extension could involve using optimization to automatically identify place fields and model them jointly.

The PTP model describes the interaction between position and theta phase as the primary factors that affect place cell activity. The interaction of these variables is specific to place cells, yet multiple variables might have similarly specific interactions that affect firing rates in other functions and regions of the spatial navigation system (31, 37-39). We hope our general statistical approach can be used to promote rigor in the study of spatial navigation and connect analyses to broader computational frameworks.

## Methods

### Parametric model of place cell activity

The model is defined by three equations, where is the position within the field and is the phase of the theta oscillation:

Spatial input equation:

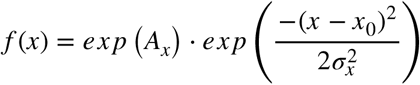

Phase modulation equation:

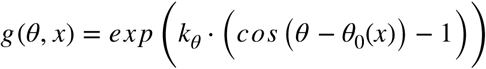

Phase precession equation:

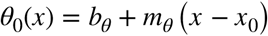

The rate is modeled as a product of the spatial input and phase modulation equations:

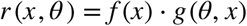

The number of spikes *k*_*t*_ occurring at position *x*_*t*_ and theta phase *θ*_*t*_ over interval *dt* is modeled as a Poisson probability distribution with mean *λ*_*t*_=*dt* · *r*(*x*_*t*_,*θ*_*t*_):

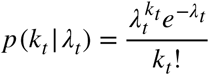

The likelihood that a model produced the data was computed as the product of probability over time:

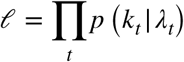

Our model explained the spiking activity of the majority of place fields better than simpler iterations of the model (Figure S3). Code, demonstrations and example data for the PTP model can be found at https://github.com/kmcclain001/ptpModel.

### Data

Spiking and local field potential were recorded from dorsal CA1 region of the hippocampus of rats as they traversed linear tracks (as described in Tingley and Buzsáki 2018). Datasets were curated for each place field by selecting time points while the animal was in each place field. The model was fit for each field using the data recorded at those time points. The inputs to the model consist of 4 time-series variables that are interpolated to the sampling rate of the local field potential (1250 Hz): 1) the position of the subject within the place field, 2) phase of the theta oscillation, 3) speed of the subject within the place field (only used in explicit speed models), 4) binary spike or no spike for each time point.

#### 1 Position

Raw position was measured as described in Tingley & Buzsaki 2018. The position on the track was linearized based on the occupancy in 2D (code included). Trials were partitioned by the starting point and running direction of the subject. Place fields were defined only within trials from a single partition. Linearized position was smoothed using a Gaussian convolution kernel and interpolated cubically to 1250Hz. Position within each place field was normalized on a 0-1 scale.

#### 2 Running speed

Speed of the rat was computed from the raw position measurements as the Euclidean distance in 2D position between frames. Speed was smoothed with a Gaussian convolution kernel and cubically interpolated to 1250Hz.

#### 3 Theta phase

To extract the theta oscillation, the local field potential was filtered using a 4th order 4-15Hz bandpass Butterworth filter. Due to speed-dependent asymmetry in the theta oscillation waveform, the phase within each cycle was defined by the latency between peaks in the signal and linearly interpolated from 0 to 2pi between consecutive peaks.

### Model fitting

Models were fit to data from each place field independently. Each time point corresponded to a datapoint with a position, theta phase, running speed and spike/no spike value. For each fit, parameters were estimated using a training dataset. A multi-start fitting procedure was used with five randomly chosen initial points to mitigate the effects of local minima in the optimization. The fmincon function in matlab was used to perform the optimization, constrained by reasonable parameter ranges (exact values can be found in code). If the parameter estimates did not converge 5 times, a field was discarded, which was the case for 80 fields.

### Parameter estimation

To assess the stability of the parameter estimates for each field, the model was fit 10 times. For each fit 90% of the data points for that field were randomly chosen to make the training dataset (Figure S1). The repeated fitting provided a distribution of parameter estimates for each field (Figure S2). The median value for each parameter was chosen as the estimate for each field.

### Model comparison

To compare the performance of competing models, Monte-Carlo cross-validation with averaging was used (40). Data were split 10 times and each model was cross-validated by fitting on 75% of the data, then testing on the remaining 25%. The mean log-likelihood for each model was computed and the model with the highest log-likelihood was chosen. Cross-validation allowed us to make a valid comparison across models with different numbers of parameters

### Neuron classification

Waveforms were clustered as described in Tingley and Buzsaki 2018. Putative cells types for each cluster were identified by four factors: firing rate, integral of the second half of the mean waveform, and the rising slope and falling slope of the autocorrelogram fit with a double exponential function. These four features were grouped using k-means clustering with 15 clusters. These clusters were merged manually into putative interneurons and putative pyramidal cells.

### Place field identification

Place fields were identified based on the firing rate of pyramidal cells (3). The mean firing rate as a function of position was computed for each cell in each trial condition. Regions on the track where the firing rate was above 20% of the peak were isolated (36). The length of these regions had to be longer than ∼1/15th the length of the track and smaller than 5/8ths the length of the track. The place cell also had to spike at least once while the subject was in the field on at least 4/5ths of trials.

## Acknowledgements

The authors would like to thank Cristina Savin and Daniel Levenstein for input on the development and implementation of the model and, the NIH training grant for computational neuroscience T90DA043219 for funding.

## Supplementary Information

**Figure S1:**
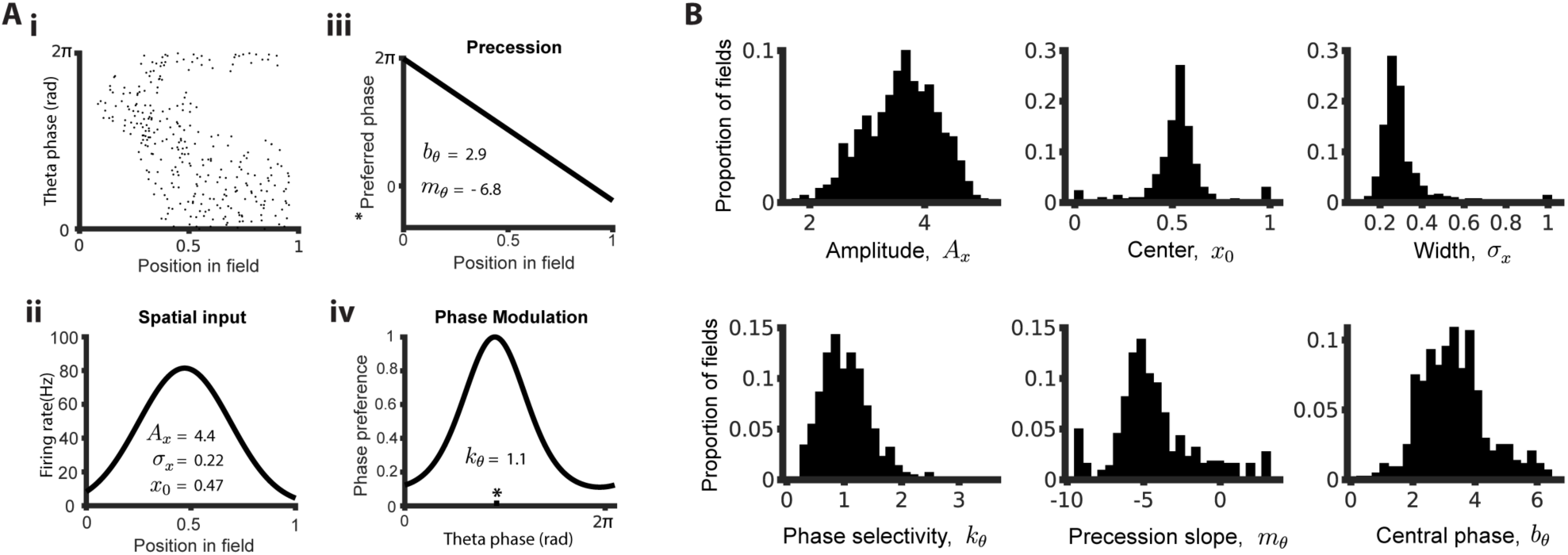
Fitting PTP model to place cell data. E) Model fit to example place field

i. Theta phase vs. position of each spike in field for real place field.
ii. Spatial input function vs. position and parameters estimated from fitting model to place field in i.
iii. Precession function vs. position.
iv. Phase modulation vs. theta phase. F) Distributions of parameter values estimated from fitting model to all place fields in data set.

**Figure S2:**
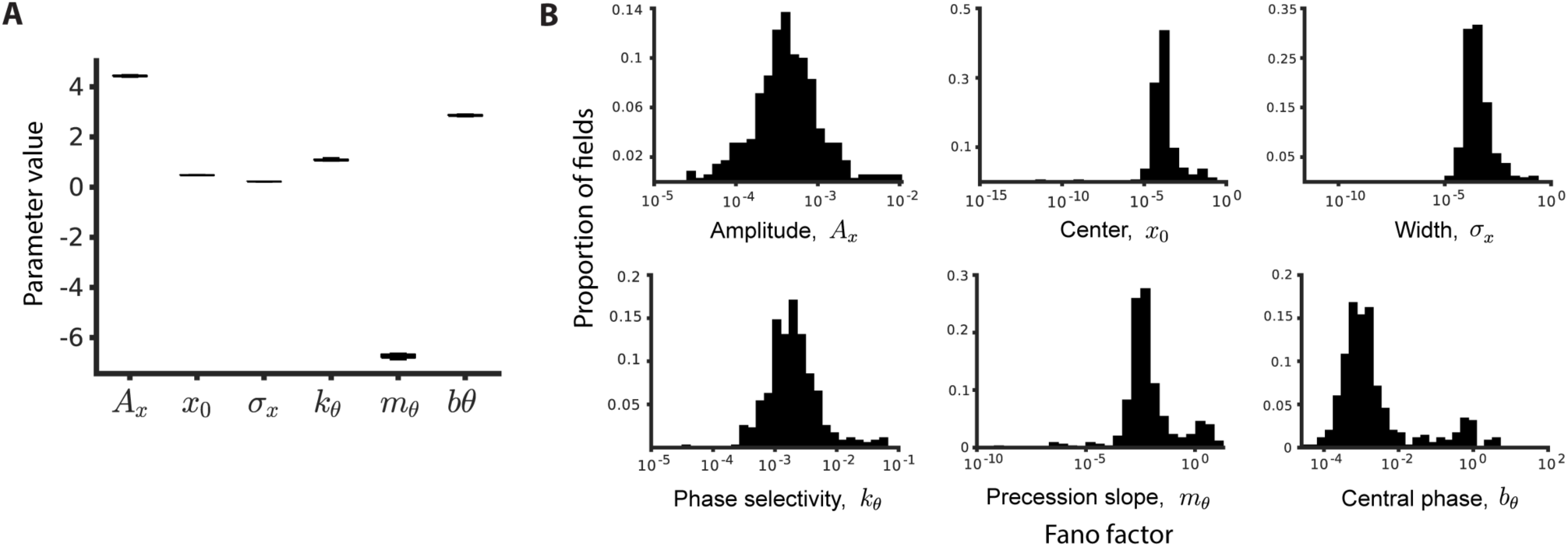
Stability of parameter estimates. A) Distribution of parameter estimates for example place field shown in Figure S1Ai. Parameters were estimated for shuffled subsets of data (see Methods). Mostly indiscernible, boxplots show median, and first and third quartiles. Whiskers represent approximately +/-2.7 standard deviations from median. B) Distribution of Fano factors for each parameter estimate across all place fields in dataset. Fano factors shown on log scale.

**Figure S3:**
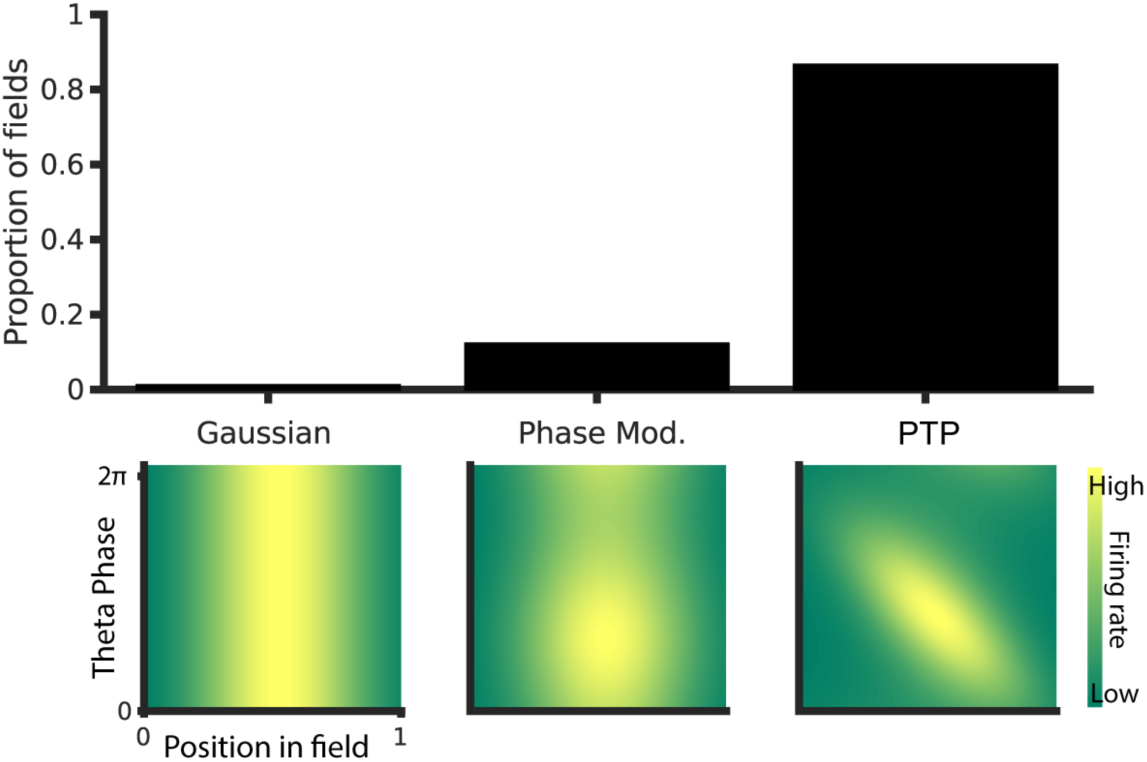
Basic model fit comparison. Performance of two simpler variants of PTP model were compared with that of the PTP model:

- Left, the simplest model was a Gaussian function of position with no theta-phase dependence.
- Middle, the next level of complexity was a stationary theta-phase modulation scaling the Gaussian, but with no phase precession.
- Right, the full PTP model as described in Figure 1. The variants are depicted under the corresponding section of the bar graph. The bar graph indicates the proportion of place fields in the data set preferring each model variant. The PTP model explains the place field data better than either of the simpler variants for the majority of place fields in the dataset.

**Figure S4:**
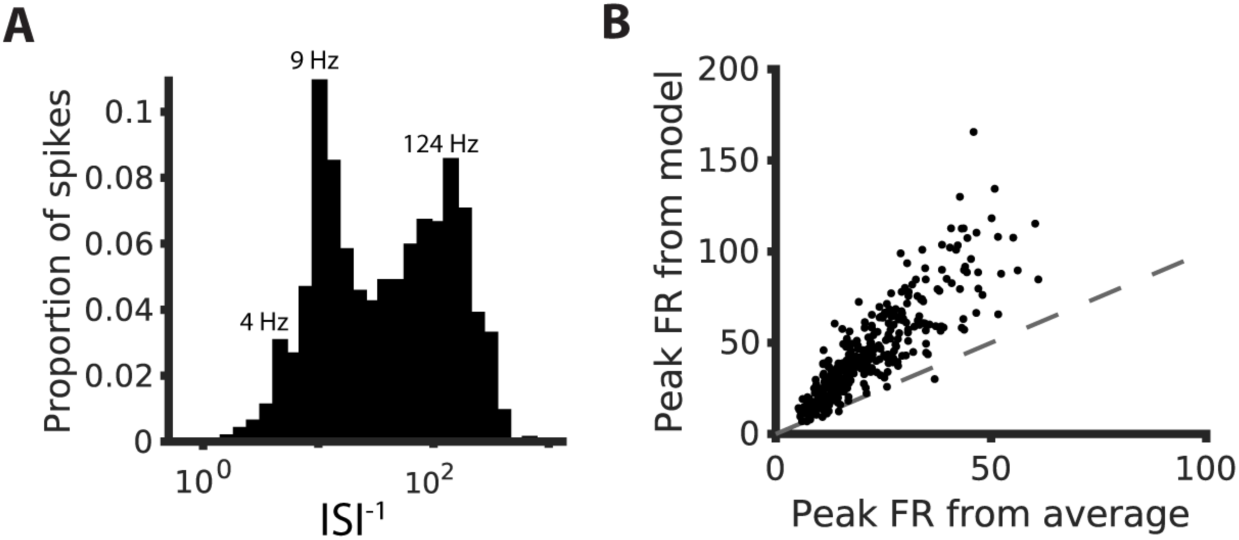
Estimating peak firing rate of place cells. A) Distribution of within-place-field inverse inter-spike intervals (ISI) across all place cells in dataset. The inverse ISI is used as an estimate of the instantaneous firing rate (41). Labeled values (4, 9, 124 Hz) correspond to lower edges of peak histogram bins. B) Peak firing rate predicted by PTP model vs. peak firing rate measured from trial-averaged firing rate for each place field.

**Figure S5:**
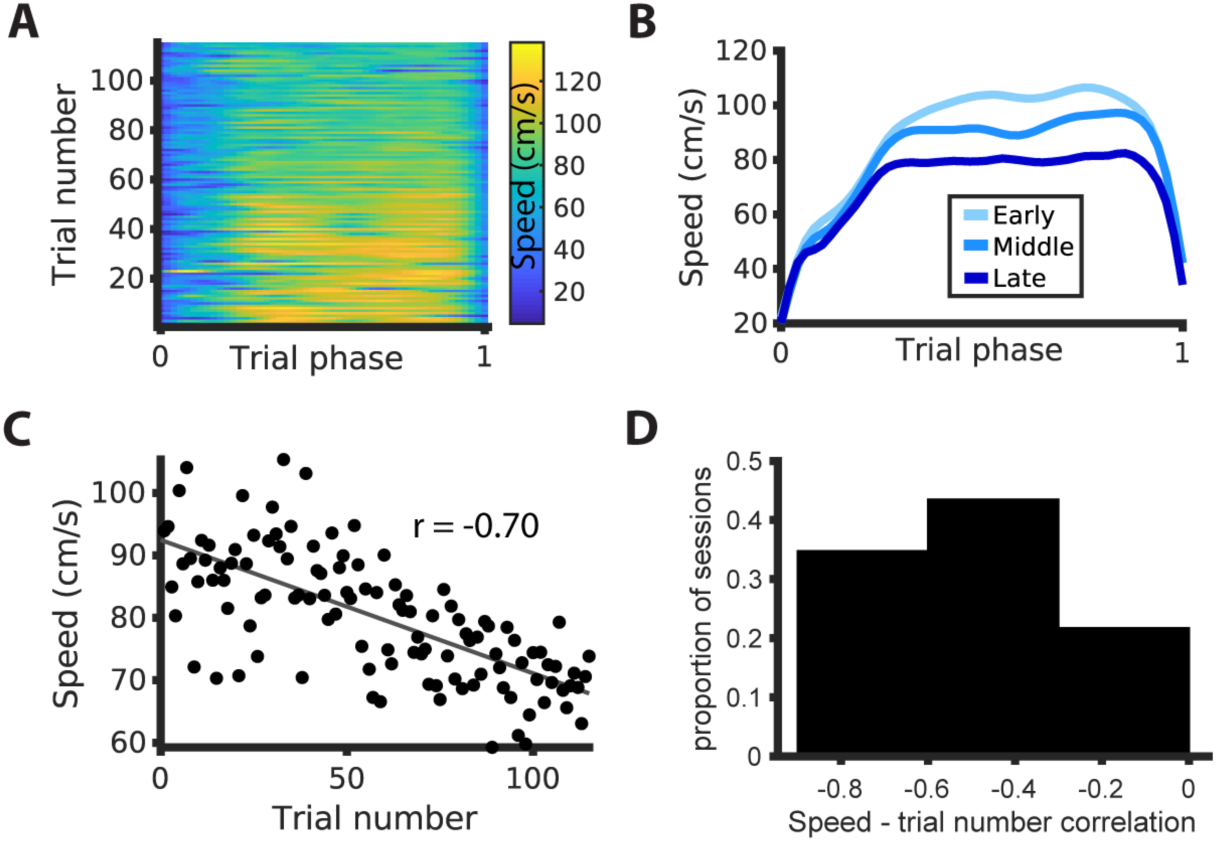
Negative correlation between running speed and trial number. A) Running speed behavior across example session. Trial lengths are normalized to a 0-1 trial phase for facility of inter-trial comparison. B) Average speed vs. trial phase for early, middle and late thirds of trials in session from A). C) Average speed within trial vs. trial number for session in A). Linear regression shown in gray. Pearson correlation coefficient between speed and trial number shown (r = −0.70). D) Distribution of Pearson correlation coefficients for speed and trial number across all behavioral sessions in dataset.

**Figure S6:**
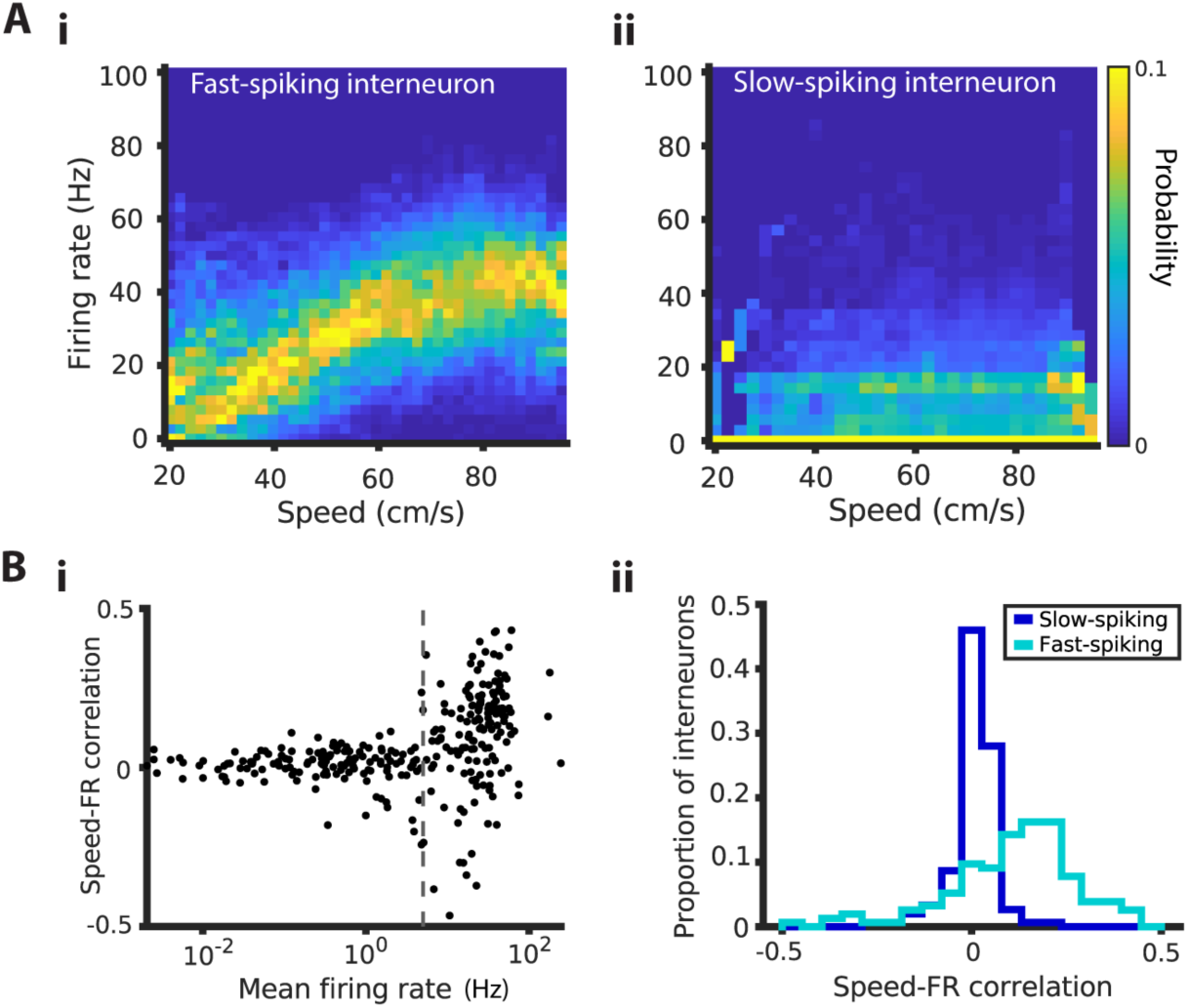
Speed dependence of firing rate in interneurons. A) Firing rate vs. speed for example interneurons: Color indicates empirical firing rate probability conditioned on running speed

i. Putative fast-spiking,
ii. Putative slow-spiking B) Speed modulation interneuron subtype specific

i. Pearson correlation coefficient between speed and firing rate vs. mean firing rate for all putative interneurons in dataset. Gray line indicates firing rate threshold (5 Hz) between putative fast-spiking and slow-spiking interneurons.
ii. Distributions of correlation coefficients for each putative interneuron subtype. Note strong positive correlation between speed and firing rate for fast firing interneurons but not for slow firing interneurons. There have been similar findings for parvalbumin- and somatostatin-expressing interneurons in entorhinal cortex (42)

